# Co-evolution networks of HIV/HCV are modular with direct association to structure and function

**DOI:** 10.1101/307033

**Authors:** Ahmed Abdul Quadeer, David Morales-Jimenez, Matthew R. McKay

## Abstract

Mutational correlation patterns found in population-level sequence data for the Human Immunodeficiency Virus (HIV) and the Hepatitis C Virus (HCV) have been demonstrated to be informative of viral fitness. Such patterns can be seen as footprints of the intrinsic functional constraints placed on viral evolution under diverse selective pressures. Here, considering multiple HIV and HCV proteins, we demonstrate that these mutational correlations encode a modular co-evolutionary structure that is tightly linked to the structural and functional properties of the respective proteins. Specifically, by introducing a robust statistical method based on sparse principal component analysis, we identify near-disjoint sets of collectively-correlated residues (sectors) having mostly a one-to-one association to largely distinct structural or functional domains. This suggests that the distinct phenotypic properties of HIV/HCV proteins often give rise to quasi-independent modes of evolution, with each mode involving a sparse and localized network of mutational interactions. Moreover, individual inferred sectors of HIV are shown to carry immunological significance, providing insight for guiding targeted vaccine strategies.

**Author summary:** HIV and HCV cause devastating infectious diseases for which no functional vaccine exists. A key problem is that while immune cells may induce individual mutations that compromise viral fitness, this is typically restored through other “compensatory” mutations, leading to immune escape. These compensatory pathways are complicated and remain poorly understood. They do, however, leave co-evolutionary markers which may be inferred from measured sequence data. Here, by introducing a new robust statistical method, we demonstrated that the compensatory networks employed by both viruses exhibit a remarkably simple decomposition involving small and near-distinct groups of protein residues, with most groups having a clear association to biological function or structure. This provides insights that can be harnessed for the purpose of vaccine design.

## Introduction

HIV and HCV are the cause of devastating infectious diseases that continue to wreak havoc worldwide. Both viruses are highly variable, possessing an extraordinary ability to tolerate mutations while remaining functionally fit. While individual residue mutations may be deleterious, these are often compensated by changes elsewhere in the protein which restore fitness [1,2]. These interacting residues form compensatory networks which provide mutational escape pathways against immune-mediated defense mechanisms, presenting a major challenge for the design of effective vaccines [3].

The compensatory interaction networks that exist for HIV and HCV—and for other viruses more generally—are complicated and far from being well understood. Resolving these by experimentation is difficult, due largely to the overwhelming number of possible mutational patterns which must be examined. An alternative approach is to employ computational methods to study the statistical properties of sequence data, under the basic premise that the residue interactions which mediate viral fitness manifest as observable mutational correlation patterns. For HIV, recent analytical, numerical and experimental studies [4–7] provide support for this premise, indicating that these patterns may be seen as population-averaged evolutionary “footprints” of viral escape during the host-pathogen combat in individual patients. This idea has been applied to propose quantitative fitness landscapes for both HIV [8,9] and HCV [10] which are predictive of relative viral fitness, as verified through experimentation and clinical data

Fitness is a broad concept that is ultimately mediated through underlying biochemical activity. For both HIV and HCV, experimental efforts have provided increased biochemical understanding of the constituent proteins, leading in particular to the discovery of small and often distinct groups of residues with functional or structural specificity (Table 1 and S1 File). These include, for example, sets of protein residues lying at important structural interfaces, those involved with key virus-host protein-protein interactions, or those found experimentally to directly affect functiona efficacy. An important open question is how these biochemically important groups relate to the interaction networks formed during viral evolution.

**Table 1.**
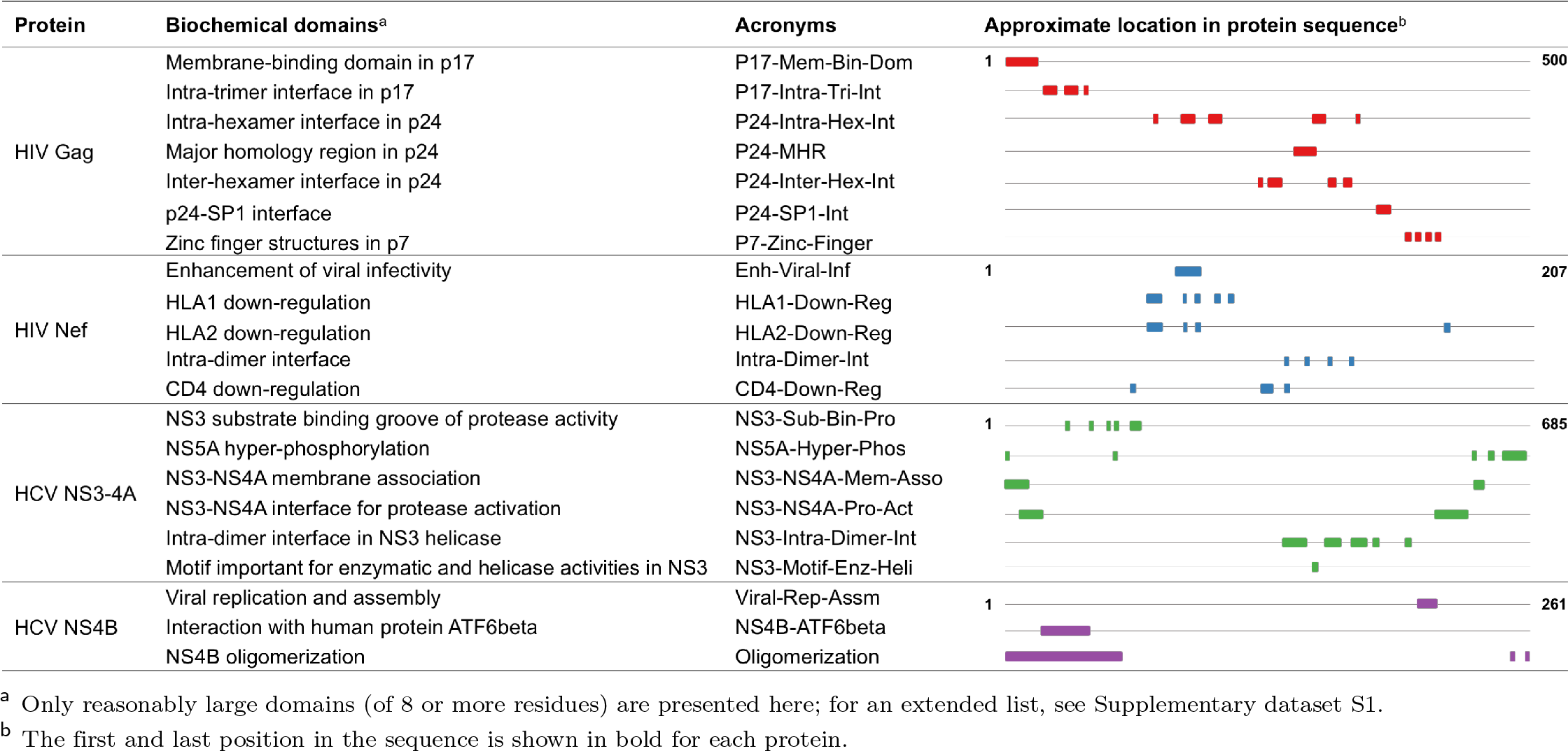
Experimentally identified biochemical domains of the viral proteins under study.

Some insights have been provided for a specific protein of HIV [11] and HCV [12]. The main objective of these investigations was to identify potential groups of co-evolving residues (referred to as “sectors”) which may be most susceptible to immune targeting. In each case, a sector with potential immunological vulnerability was inferred and this was found to embrace some residues with functional or structural significance It is noteworthy that the employed inference methods were designed to produce strictly non-overlapping groups of co-evolving residues, which may hinder identification of inherent co-evolutionary structure and associated biochemical interpretations.

In a parallel line of work, computational methods have been used to understand co-evolution networks for various protein families, with compelling results (reviewed in [13]). Notably, for the family of S1A serine proteases [14], a correlation-based method termed statistical coupling analysis (SCA) uncovered a striking modular co-evolutionary structure comprising a small number of near-independent groups of co-evolving residues (again referred to as sectors), each bearing a clear and distinct biochemical association in addition to other qualitative properties. Sectors have also been obtained for other protein families using SCA, and the functional relevance of these has been confirmed through experimentation [14–17]. A natural question is to what extent such modular, biochemically-linked co-evolutionary organization exists for the viral proteins of HIV and HCV? This is not obvious, particularly when one considers the evolutionary dynamics of these viruses, which are complex and very distinct to those of protein families. Specifically, they involve greatly accelerated mutation rates, and are shaped by effects including intrinsic fitness, host-specific but population-diverse immunity, recombination, reversion, genetic drift, etc. The sampling process is also complicated and subject to potential biases, making inference of co-evolutionary structure difficult.

In this paper, considering multiple proteins of HIV and HCV, we identify in each case a sparse and modular co-evolutionary structure, involving near-independent sectors. This is established by introducing a statistical method, which we refer to as “robust co-evolutionary analysis (RoCA)”, that learns the inherent co-evolutionary structure by providing resilience to the statistical noise caused by limited data. Strikingly, the sectors are shown to distinctly associate with *often unique* functional or structural domains of the respective viral protein, indicating clear and well-resolved linkages between the evolutionary dynamics of HIV and HCV viral proteins and their underlying biochemical properties. Our results suggest that distinct functional or structural domains associated with each of the viral proteins give rise to quasi-independent modes of evolution. This, in turn, points to the existence of simplified networks of sparse interactions used by both HIV and HCV to facilitate immune escape, with these networks being quite localized with respect to specific biochemical domains. The insights provided by the inferred sectors also carry potential importance from the viewpoint of immunology and vaccine design, which we demonstrate for a specific protein of HIV.

## Results

### Modular and sparse co-evolutionary structure of HIV/HCV proteins

By employing available sequence data, we investigated the co-evolutionary interaction networks for various viral proteins. Specifically, we considered the Gag and Nef proteins of HIV and the NS3-4A and NS4B proteins of HCV. Our proposed approach, RoCA, resolves co-evolutionary structures by applying a spectral analysis to the mutational (Pearson) correlation matrices and identifying inherent structure embedded within the principal components (PCs). The RoCA algorithm is designed to be robust against statistical noise, which is a significant issue since the number of available sequences for each protein is rather limited, being comparable to the protein size (Fig 1). After an initial screening step to isolate PCs which carry correlation information from those which are supposedly dominated by statistical noise, as a key component of RoCA, we developed a suitably adapted version of a statistical technique [18], which provides robust estimates of the PCs. To summarize, this procedure involves: (i) a thresholding step that distinguishes, for each PC, the significantly correlated residues from those residues whose correlations are consistent with statistical noise, and (ii) an iterative procedure that tries to robustly estimate the correlation structure between the selected residues across different PCs (see Fig 1 and Materials and methods for details). Based on the resulting PCs, the RoCA algorithm directly infers co-evolutionary sectors, representing groups of residues whose mutations are collectively coupled. Importantly, other than applying a suitable data-driven thresholding step to remove statistical noise, the method makes no structural imposition on the inferred sectors, and it is therefore designed to reveal inherent co-evolutionary networks as reflected by the data.

**Fig 1.**
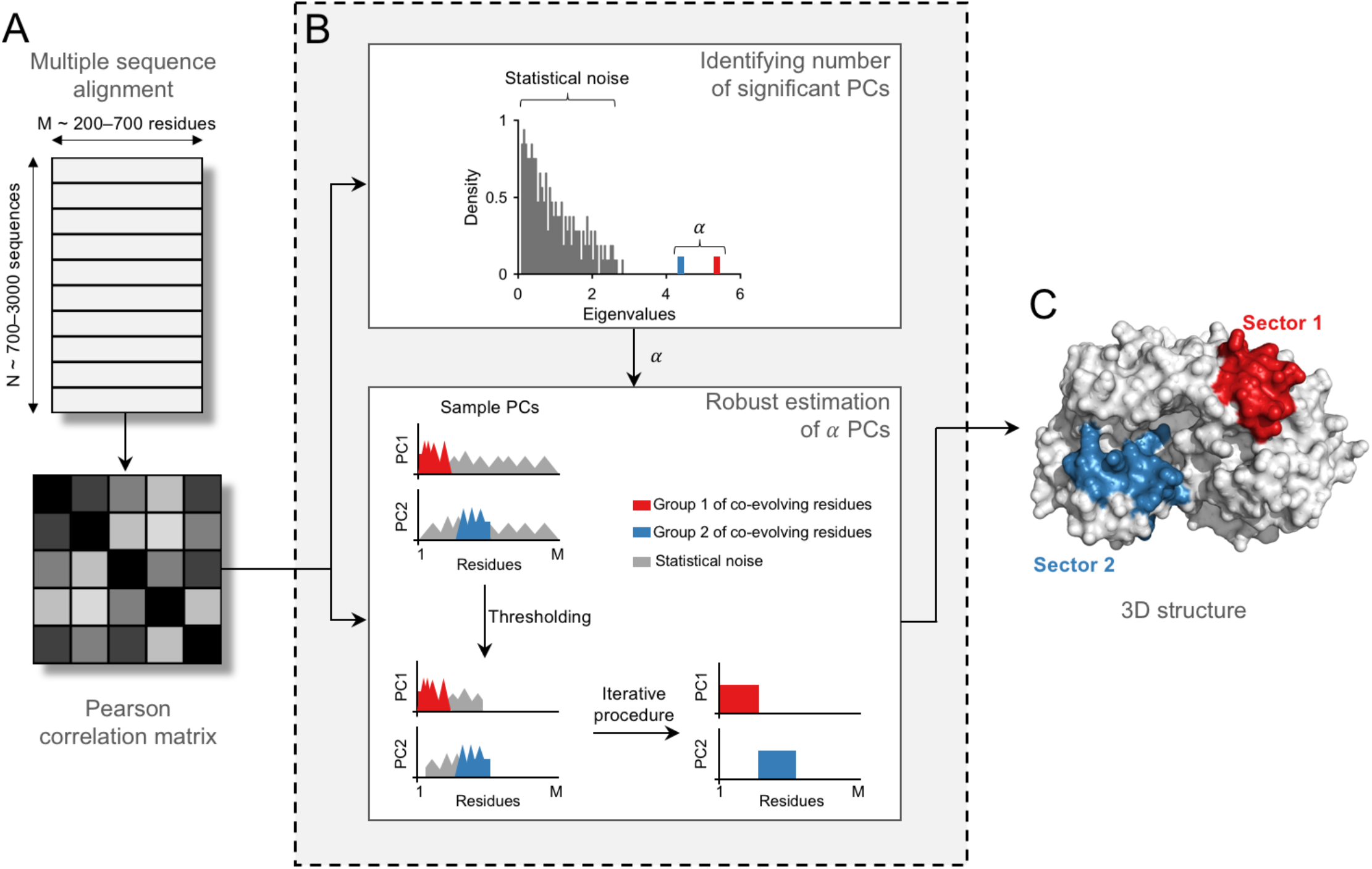
Inferring co-evolutionary structure using the RoCA method. Illustration of the RoCA algorithm for a simple toy model involving two non-overlapping sectors of co-evolving residues. (A) Data pre-processing that involves computation of the mutational Pearson correlation matrix from a multiple sequence alignment. (B) (*Top panel*) A spectral analysis on the correlation matrix is performed to distinguish true correlations, encoded in the dominant spectral modes (shown here in red and blue colors), from those which seemingly reflect statistical noise. The observed eigenvalue spectrum is reminiscent of that generally observed in *spiked correlation models* [19], which includes a bulk of small eigenvalues representing largely statistical noise and a few big eigenvalues (referred to as spikes) representing the true underlying correlations. (*Bottom panel*) The dominant PCs are estimated to identify the co-evolutionary structure using the proposed robust method. This involves an intelligent data-driven thresholding step based on random matrix theory to identify the set of all correlated residues (those present in both sectors) from statistical noise, followed by an iterative procedure to determine the correlated residues associated with each PC from the set of all correlated residues. Based on the resulting PCs, the groups of co-evolving residues (sectors) are accurately identified. Note that these groups are not necessarily contiguous in the primary sequence, as assumed in this toy model construction. (C) Sectors, inferred using the robustly estimated PCs, are generally closely placed in the 3D structure.

For each viral protein, the RoCA method identified a small number of sectors (Fig 2A and S1 Fig) which together embraced a rather sparse set of residues (i.e., between 35%-60% of the protein; see S2 File for the complete list). In some cases the sector residues were localized in the primary sequence, while in others they were quite well spread (Fig 2B and S1 Fig). Importantly, while each sector was identified from a distinct PC, they were found to be largely disjoint (Fig 2A and S1 Fig). This suggests that the co-evolutionary structures are highly modular, with the different modules (sectors) being nearly uncorrelated to each other. In fact, further statistical tests demonstrated that the inferred sectors are nearly independent (Fig 2C and S1 Fig).

**Fig 2.**
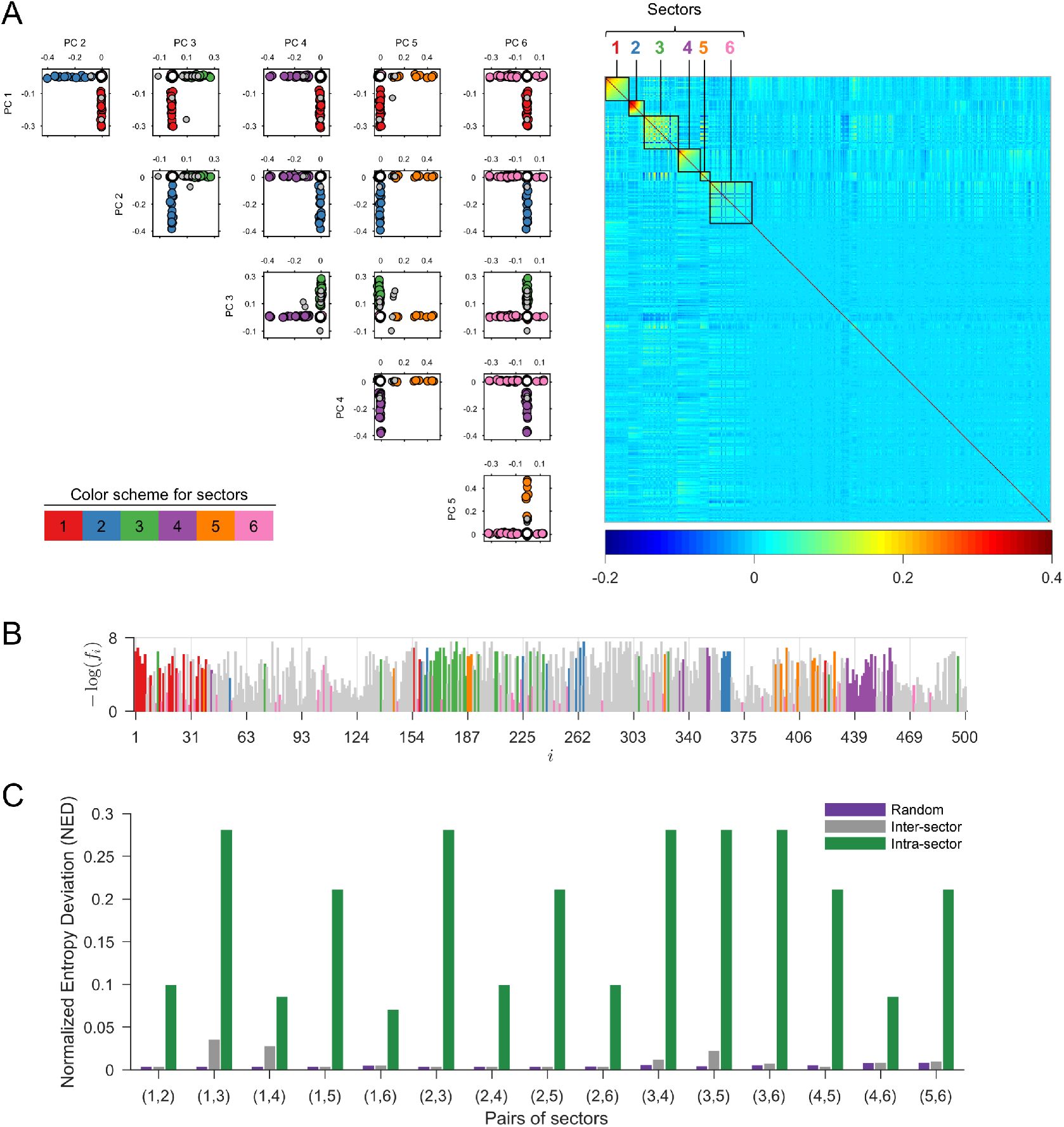
Co-evolutionary sectors revealed by RoCA for HIV Gag. (A) Biplots of the robustly estimated PCs that are used to form RoCA sectors. The sector residues are represented by circles according to the specified color scheme, while overlapping residues (belonging to more than one sector) and independent (non-sector) residues are represented as gray and white circles, respectively. The heat map of the cleaned correlation matrix (Materials and methods), with rows and columns ordered according to the residues in the RoCA sectors, shows that the sectors are notably sparse and uncorrelated to each other. (B) Location of RoCA sector residues in the primary structure of HIV Gag. The sector residues are colored according to the specifications in (A) while remaining residues are shown in gray color. The vertical axis of each plot shows the negative log-frequency of mutation for each residue *i*. (C) Statistical independence of sectors using the normalized entropy deviation (NED) metric. It is a non-negative measure which is zero if two sectors are independent, while taking a larger value as the sectors become more dependent (see Materials and methods for details). For each possible pair of sectors, the inter-sector NED is very small and generally close to the randomized case, while being substantially lower than the maximum intra-sector NED of any individual sector in the considered pair, reflecting that sectors are nearly independent. Corresponding results for the other three proteins are presented in S1 Fig.

This identified modular co-evolutionary structure is in fact reminiscent of ‘community structure’ that has been observed in numerous complex networks, e.g., metabolic, webpage, and social networks [20]. In such applications, the identified modules or communities have been shown to represent dense sub-networks which perform different functions with some degree of autonomy. For our co-evolutionary sectors, in line with previous studies on the fitness landscape of HIV [6–8] and HCV [10], they appear to represent groups of epistatically-linked residues which work together to restore or maintain viral fitness when subjected to strong selective pressures during evolution (e.g., as a consequence of immune pressure). In light of this, one anticipates that the co-evolutionary sectors should afford an even deeper interpretation in terms of the underlying biochemical properties of the viral proteins, which fundamentally mediate viral fitness.

### Modularity in HIV/HCV is tightly coupled to biochemical domains

To explore potential correspondences between the identified RoCA sectors and basic biochemical properties, we compiled information determined by experimental studies for each of the viral proteins. This consists of residue groups having prescribed functional or structural specificity (see Table 1; also S1 File for a more extensive list including small groups). These groups, which are seen to occupy sparse and largely distinct regions of the primary structure (Table 1), are collectively referred to as “biochemical domains”. (This should not be confused with the term “domain”, as classically used for a folding unit in structural biology and biochemistry.) For each viral protein, structural domains were defined based on spatial proximity of residues in the available protein crystal structure; they include, for example, residues which lie on critical interfaces needed to form stable viral complexes, or those involved in essential virus-host protein-protein interactions. Functional domains, on the other hand, were typically identified using site-directed mutagenesis or truncation experiments, and they include groups of residues found to have a direct influence on the efficacy of specific protein functions. It is important to note, however, that while structural domains are typically clearly specified, functional domains are expected to be less so, due to experimental limitations. Results reported based on truncation experiments, for example, may comprise false positives due to the coarse nature of the experimental procedure.

Despite potential limitations of the compiled biochemical domains, contrasting these domains with the RoCA sectors (for all four viral proteins) revealed a striking pattern, with most sectors showing a clear and highly significant association to a *unique* biochemical domain (Fig 3). This is most marked for the HIV Gag protein, where there is a one-to-one correspondence. These observations carry important evolutionary insights. Not only are the co-evolutionary networks of both HIV and HCV proteins modular, but the modules (sectors) seem to be intimately connected to distinct biological phenotype. Our results suggest that the fundamental structural or functional domains of these viral proteins spawn quasi-independent co-evolutionary modes, each involving a simplified sparse network of largely localized mutational interactions. The observed phenomena is seemingly a natural manifestation of immune targeting against residues within the biochemical domains, since mutations at these residues likely lead to structural instability or functional degradation, necessitating the formation of compensatory mutations to restore fitness and facilitate immune escape.

**Fig 3.**
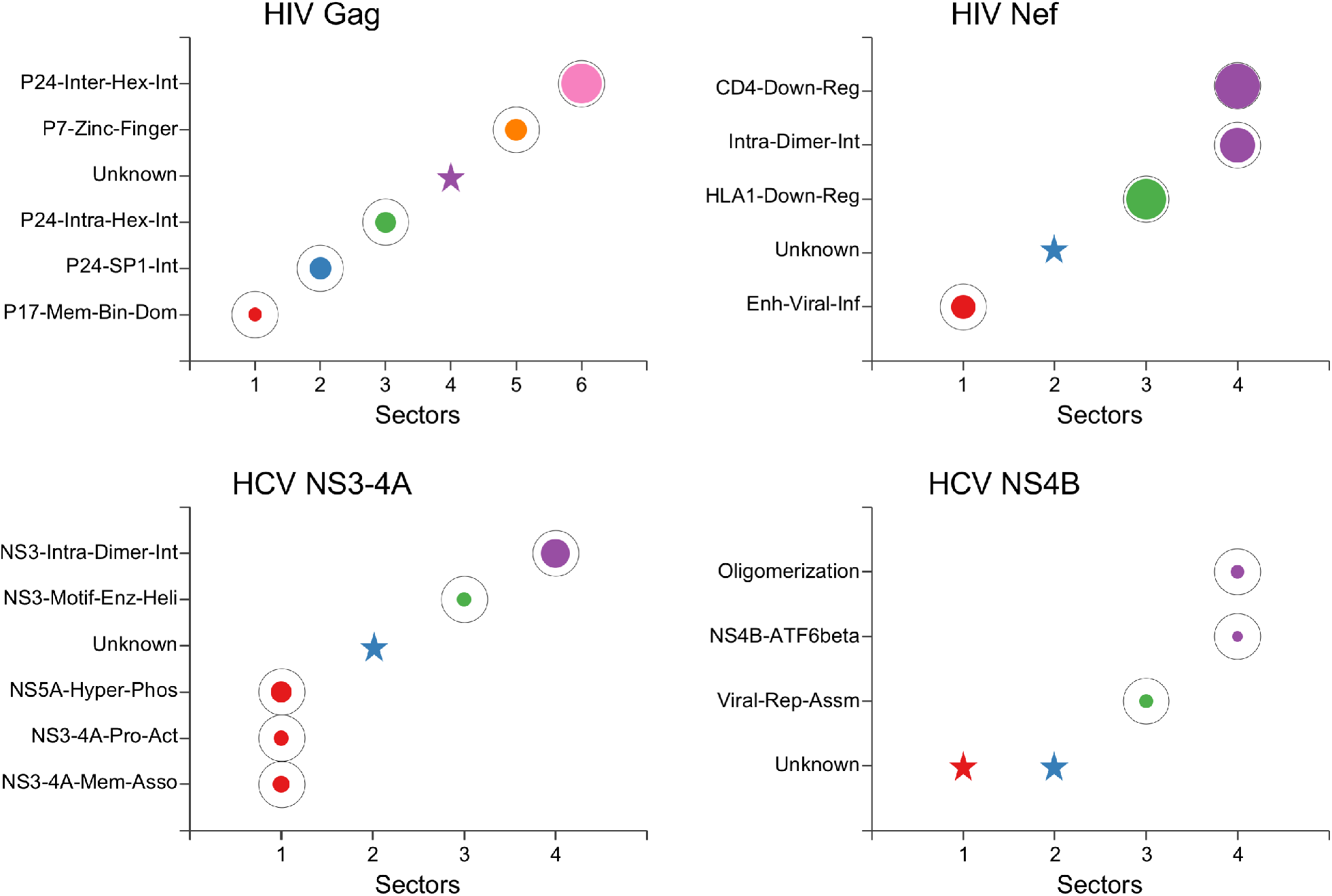
Individual associations of RoCA sectors with the biochemical domains of the studied viral proteins. The sectors are colored according to the scheme in Fig 2A. The area of each bubble reflects the statistical significance of the associated result, measured as –1/ log_10_ *P*, where *P* is the *P*-value computed using Fisher’s exact test, and the black circles indicate the conventional threshold of statistical significance, *P* = 0.05; any *P*-value lower than that (bubble inside the black circle) is considered statistically significant. The star symbols indicate those RoCA sectors with unknown biochemical significance. Note that the involved structural interfaces were defined based on a contact distance of less than *d* = 7Å between the alpha-carbon atoms. Similar qualitative results are obtained for *d* = 8Å or *d* = 9Å (S2 Fig).

### Co-evolutionary structure identified with previous sectoring methods

We investigated whether our main findings could also be revealed by other (previous) sectoring methods. We first investigated a method which we proposed previously based on classical PCA [12] (a slight variant of the approach in [11]), which sought to identify groups of collectively-correlated viral residues which may be susceptible to immune targeting. An important feature of the algorithm was the imposition of a structural constraint in the inferred sectors, enforced to be disjoint [12], which may compromise its ability to infer natural co-evolutionary structure (S1 Text). Despite the imposed constraints, the sectors produced by this method for the studied viral proteins tended to be larger than the RoCA sectors (S3 Fig), and they collectively embraced a larger set of residues (covering 40%–80% of the protein). In many cases, they included a mix of residues from multiple RoCA sectors (Fig 4A), a fact that was also reflected in the biochemical associations of the sectors, where much of the resolved (unique) ssector/domain associations shown by RoCA (Fig 3) were indeed no longer revealed (S3 Fig). We found that these key differences were also attributed to the sensitivity of the approach to sampling noise (limited sequence data), as reflected by the noisy and significantly overlapping principal components (Fig 4B). This was corroborated with a ground-truth simulation study, through which the ability to infer co-evolutionary structure was tested in synthetic model constructions (S2 Text). The RoCA method resolved all the individual (true) sectors with high accuracy, whereas our previous method [12] inferred comparatively large sectors, which often included false positives and merged residues from different true sectors (S3 Fig).

**Fig 4.**
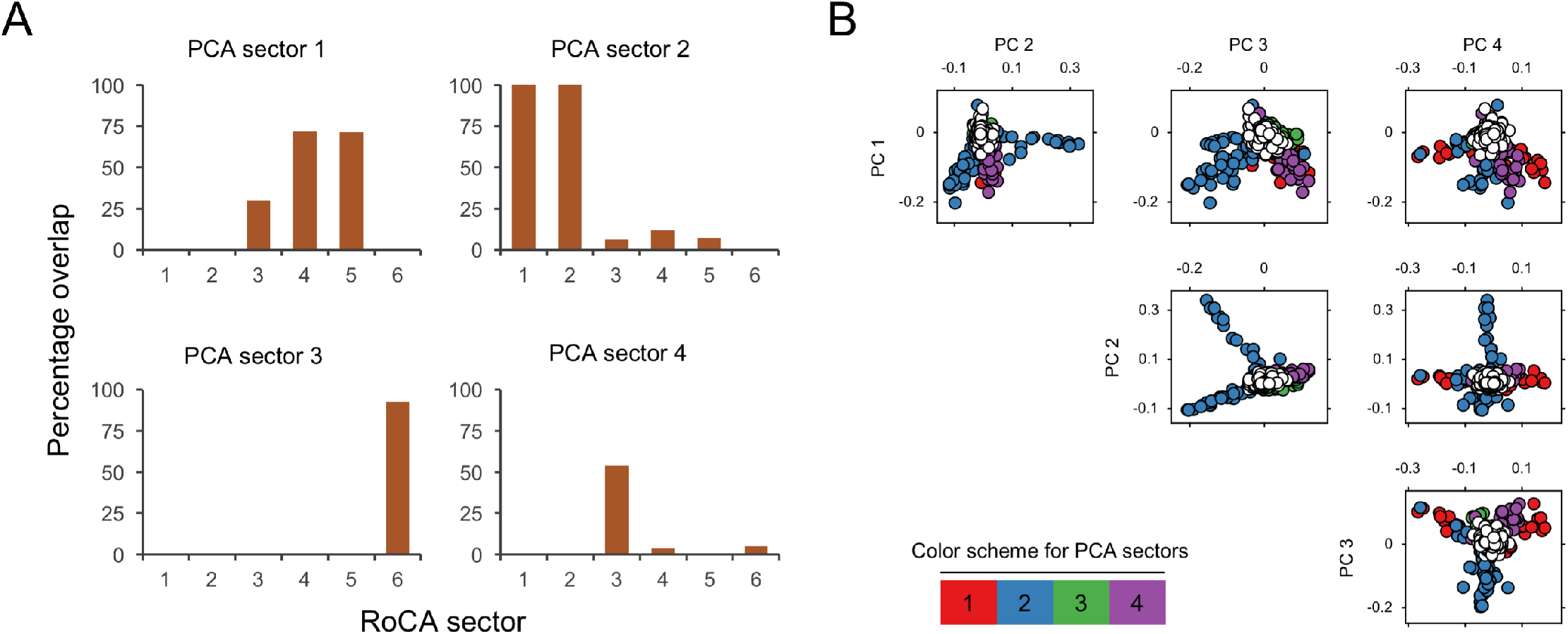
Sectors revealed by the PCA-based method [12] for HIV Gag. (A) Bar plots showing the merging of multiple RoCA sectors in the sectors revealed by the method in [12] (only the first four are shown). The vertical axis of each plot shows the percentage of residues within the different RoCA sectors that fall into the prescribed sector. (B) Biplots of the PCs which are post-processed to form sectors in [12]. The sector residues are represented by circles according to the specified color scheme, while independent (non-sector) residues are represented as white circles. The PCs can be seen to be severely affected by statistical noise. Note the substantial overlap in the support (relevant entries) of the PCs; such overlap is however not present in the formed sectors, as the method in [12] applies heuristic post-processing steps to enforce disjoint sectors. Corresponding results for the other three proteins are presented in S4 Fig.

Our main findings were neither established with other co-evolutionary methods, which tended to return very different results to RoCA, and generally revealed little biochemical association for the studied viral proteins (S5 Fig). Most notable is the benchmark SCA method [14] which has shown much success in resolving co-evolutionary structure for certain protein families [15,21] (S5 Fig). Aside from the noise sensitivities shared by both SCA and classical PCA-based methods (S5 Fig), the surprising disparity in this case appears due to the weighted covariance construction employed by SCA (as opposed to the Pearson correlation) which, while being suited to the analysis of certain protein families data [14–17], does not seem suitable for identifying the co-evolutionary structure in the considered HIV and HCV proteins (see S3 Text for details).

### Detailed analysis of the biochemical associations of the inferred sectors

In the following, we provide details on the biochemical associations of the identified RoCA sectors for each of the four viral proteins.

### HIV Gag

Gag poly-protein encodes for the matrix (p17), capsid (p24), spacer peptide 1 (SP1), nucleocapsid (p7), spacer peptide 2 (SP2), and p6 proteins. Being a core structural poly-protein of HIV, the experimentally identified domains in Gag consist of critical structural interfaces involving either virus-host or virus-virus protein interactions (Table 1).

Strikingly, five of the six identified RoCA sectors were individually associated to distinct structural domains (Fig 5A). Sector 1 was enriched (52%) with N-terminal residues of p17 involved with virus-host protein interaction—binding of Gag with plasma membrane—critical for viral assembly and release [22]. The remaining sectors were associated with virus-virus protein interactions. In particular, sector 2 consisted of a large proportion of residues (56%) that form the p24-SP1 interface, which is considered to be important for viral assembly and maturation [23]. Sector 3 was dominated by the residues belonging to the capsid protein p24. The HIV capsid exists as a fullerene cone with 250 hexamers and 12 pentamers that cap the ends of the cone. The monomer-monomer interface formed in the oligomerization of p24 (in both the hexamer and pentamer structures) has been shown to be important for structural assembly of the HIV capsid [24,25]. Sector 3 was enriched with ~50% of the residues in the largely overlapping interfaces of these p24 oligomers (S1 File). In addition to these residues, the hexamer-hexamer interface in p24 has been shown to be important for proper capsid formation [26]. Sector 6 was found to comprise 36% of the residues within this interface. Sector 5 consisted of a large proportion (40%) of residues involved in a critical functional domain—two zinc finger structures separated by a basic domain—in p7, important for packaging genomic RNA [27].

**Fig 5.**
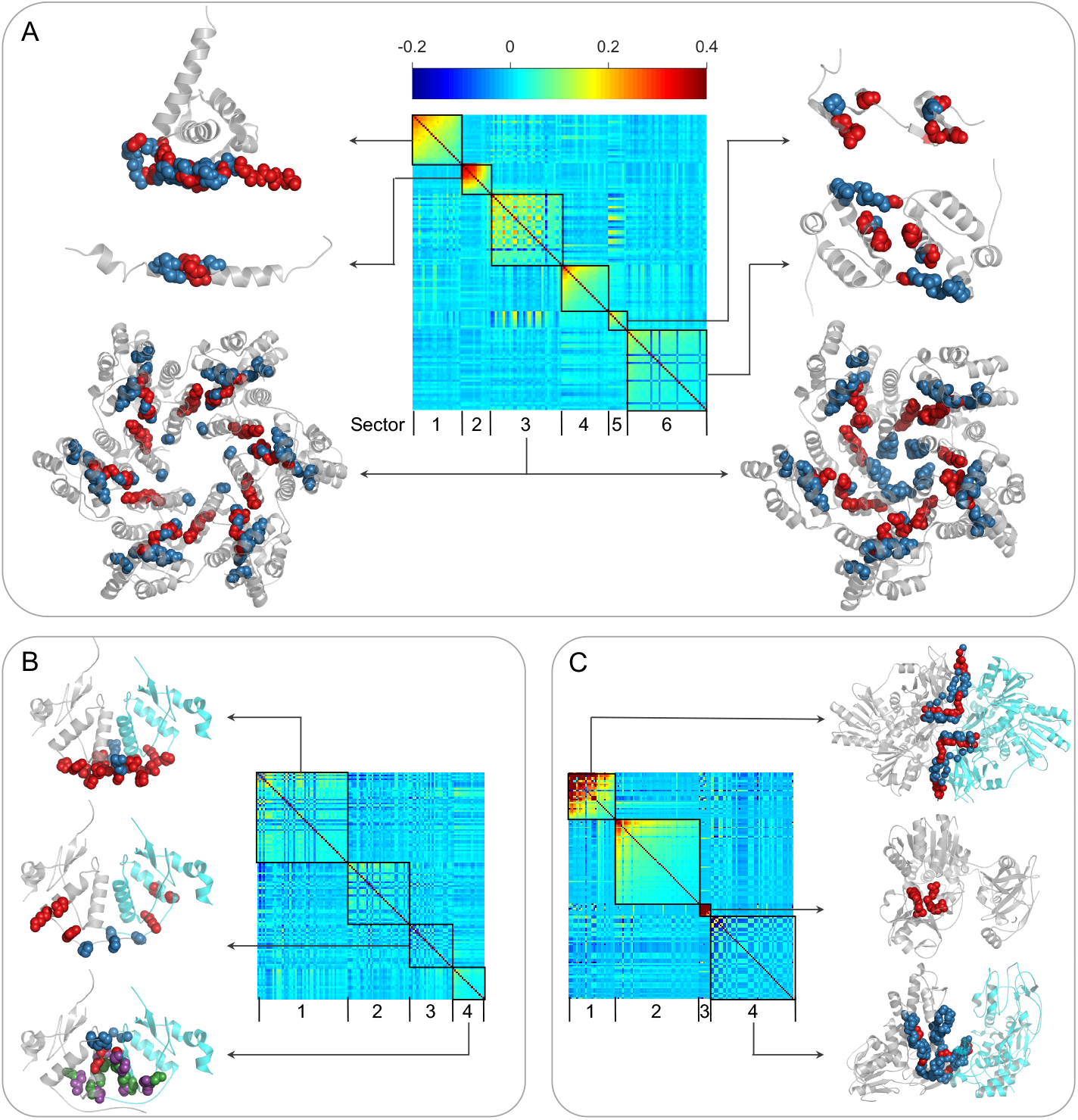
Details of the different biochemical domains of viral proteins associated with the respective inferred RoCA sectors. Sectors are shown as diagonal blocks in the heat map of cleaned correlations with rows and columns re-ordered accordingly (Materials and methods); the heat map is restricted to sector residues only. For each sector, the crystal structure of the associated domains are depicted, when available. The protein chains are shown in gray (in case of dimer structures, chains A and B are depicted in gray and cyan colors, respectively), the relevant domain residues present in the sector are represented as red spheres, and the remaining domain residues are shown as blue spheres; the side chains of all the atoms are excluded for clarity of presentation. (A) For HIV Gag, sector 1 residues are associated with the membrane-binding domain of p17 (PDB ID 2LYA); sector 2 residues with the p24-SP1 interface (PDB ID 1U57); sector 3 residues with the intra-hexamer and intra-pentamer interface of p24 (PDB ID 3GV2 and 3P05, respectively); sector 5 residues with the two zinc-finger structures of p7 (PDB ID 1MFS); and sector 6 residues with the inter-hexamer interface of p24 (PDB ID 2K0D). No crystal structure is available for the SP2-p6 interface residues that comprise sector 4. (B) For HIV Nef, relevant residues of the biochemical domains are shown on the dimer crystal structure (PDB ID 4U5W). Note that this crystal structure only includes residues 68-204 of Nef Sector 1 residues are associated with the viral infectivity enhancement function; sector 3 residues with HLA1 down-regulation function (note that four residues (62-65) of this biochemical domain cannot be shown in this crystal structure); and sector 4 residues with both the CD4 down-regulation function and oligomerization (for the latter domain, the residues present in the sector are represented as green spheres and the remaining domain residues are shown as purple spheres). (C) For HCV NS3-4A, sector 1 residues are associated with the interface between NS3 and NS4A proteins (PDB ID 4B6E) important for activation of the NS3 protease function (this crystal structure only includes NS4A residues 21-36); sector 3 residues with the motif critical for enzymatic and helicase activities of NS3 (PDB ID 4B6E; no blue sphere is visible as sector 3 comprises ah the residues in this motif); and sector 4 residues with the hehcase-hehcase interface of the NS3 dimer (PDB ID 2F55).

We found that sector 4 (indicated as a star in Fig 3) was not significantly associated with any of the large biochemical domains identified for HIV Gag (listed in Table 1). This sector comprised the complete SP2 protein and a few N-terminal residues of the p6 protein, for which little experimental information is available. To our knowledge, only five of these residues were experimentally studied previously [28,29], wherein mutations were shown to alter protein processing and abolish viral infectivity and replication. While the biochemical implications for the remaining residues in sector 4 are not known, our result suggests that they could also be important for the mentioned functions. These residues thus serve as potential candidates for further experimental studies. A similar comment applies for each of the proteins discussed below in relation to those sectors with as yet unspecified biochemical association.

### HIV Nef

Nef is an accessory protein which is a critical determinant of HIV pathogenesis and is involved in multiple important functions.

Of the four RoCA sectors revealed for Nef, two of these (sectors 1 and 3) were associated with distinct biochemical domains, while one (sector 4) was associated with two domains (Fig 5B). Specifically, sector 1 was enriched (90%) with residues in a functional domain consisting of the proline-x-x repeat, shown to be critical for the enhancement of viral infectivity [30], while sector 3 consisted of a large proportion (44%) of residues considered to be crucial in virus-host protein interactions that down-regulate the surface expression of HLA1 molecules [31]. In contrast, sector 4 comprised (i) 33% of the residues involved in the virus-host protein interaction that results in the down-regulation of CD4 surface expression [32], and (ii) 44% of the residues involved in the virus-virus protein interaction critical for Nef dimerization [33]. Although the residues in these two biochemical domains are close in the primary structure (Table 1), they are largely distinct with only one residue in common. The association of a single co-evolutionary sector with these two domains suggests, however, that they may be biochemically related, with mutations in one domain influencing the other. Interestingly, this is corroborated by a recent study [34] which shows that mutations disrupting the Nef dimer structure highly affect the CD4 down-regulation function. Further experimental work is still required to more finely resolve the dependencies between these domains. Nonetheless, the association of viral infectivity enhancement and CD4 down-regulation with distinct sectors (1 and 4) is remarkably in line with the experimentally reported dissociation of these two functions [30].

We found a single sector (sector 2) that was not associated with any known biochemical domain (Fig 3 and Fig 5B). The available crystal structure suggests that these residues are predominantly located away from the dimer interface, yet the biochemical significance of these residues remains unknown.

### HCV NS3-4A

NS3 is a large protein involved with performing serine protease and helicase functions. Based on these functions, NS3 is divided into two domains: the protease domain, consisting of N-terminal one-third protein residues, and the helicase domain, comprising the remaining C-terminal two-third protein residues. NS4A is a very short protein that functions as a co-factor for the serine protease activity of NS3.

Of the four RoCA sectors, two (sectors 3 and 4) were individually associated with distinct biochemical domains, while one (sector 1) was associated with multiple domains (Fig 5C). Specifically, the small sector 3 contained all the residues of a relatively conserved motif in the NS3 helicase domain, considered to be important for ATPase and duplex unwinding activities [35], while sector 4 comprised 31% of the residues involved in dimerization of NS3, important for helicase activity and viral replication [36]. In contrast, sector 1 was a mixture of the NS3 protease domain and NS4A residues, encompassing multiple functional domains. In particular, the N-terminal residues of the NS3 protease domain have been reported to be involved in three virus-virus protein interactions with NS4A to mediate multiple functions including: (i) activation of the NS3 protease function [37]; (ii) membrane association and assembly of a functional HCV replication complex [38]; and (iii) NS5A hyper-phosphorylation [39,40]. Sector 1 comprised 40%–52% of the residues associated with these functions. The association of a single sector with these multiple domains (Table 1) in NS3-4A suggests that they may be functionally coupled. Interestingly, this is in line with experimental studies which report the dependence of NS3-4A membrane association and NS5A hyper-phosphorylation on an active NS3 protease [38,40]. Further work is still required to more finely resolve these dependencies.

For NS3-4A, sector 2 was not associated with any known biochemical domain (Fig 3 and Fig 5C). This sector included residues that are well-distributed in the NS3 protease and helicase domains as well as in NS4A.

### HCV NS4B

NS4B is a small hydrophobic membrane protein which is involved in multiple functions including viral replication and assembly. Compared to other HCV proteins, NS4B is relatively poorly characterized and its full-length crystal structure is still not available.

Of the four RoCA sectors, two were associated with known biochemical domains (Fig 3). Sector 3 was strongly associated with a functional domain comprising the C-terminal *α*-helix 1 (H1), shown to be important for HCV RNA replication and viral assembly [41]. This sector consisted of all H1 residues, but none of the *α*-helix 2 (H2) residues. This is in agreement with [41] which reported a comparably higher impact of H1, as compared with H2, on viral replication and assembly.

Sector 4 comprised half of the residues considered to be involved in virus-host protein interaction between a basic leucine zipper (bZIP) motif at the N-terminal of NS4B and the central part of the human protein ATF6beta (activating transcription factor 6 beta) [42]. Moreover, this sector also contained residues within a region that, at a coarse level, was identified to be sufficient for NS4B oligomerization by a truncation procedure [43]. Thus, while the specific residues involved in NS4B oligomerization remain unknown, sector 4 may assist in a more accurate identification of these residues, thereby refining the coarse analysis of [43]. The bZIP motif residues completely overlap with this biochemical domain (Table 1) and thus both were associated with the same sector.

With the limited current understanding of the functional and structural characteristics of NS4B, sectors 1 and 2 could not be associated to any known biochemical domain (Fig 3). Nonetheless, we examined the predicted secondary structure of the NS4B protein [44] to gain some insight. Both sectors consisted of residues present in the central part of NS4B that contains the trans-membrane (TM) segments. Specifically, sector 1 comprised the majority of residues in TM3, while sector 2 consisted of residues in TM1 and TM2. These TM segments are considered to be important for mediating the membrane-association of NS4B [44]. However, the specific residues involved with this function are still unresolved, and therefore the corresponding association with sectors 1 or 2 could not be clearly established.

### Association of sectors with viral control and disease progression of HIV

Our main results carry potential immunological significance, which may provide useful input for vaccine design. To demonstrate this, we considered the HIV Gag protein, and contrasted the sectors with the epitope residues targeted by T cells of HIV “long-term non-progressors” (LTNP) and “rapid progressors” (RP). LTNP correspond to rare individuals who keep the virus in check without drugs, whereas RP are individuals who tend to progress to AIDS in less than 5 years (compared to the population average of 10 years [45]). Our analysis revealed that LTNP elicit immune responses strongly directed towards residues in sector 3, whereas RP elicit responses against residues in sector 2 (S1 Table and Fig 6). Recalling the sector biochemical associations (figures 3 and 5), these observations seem to promote the design of T-cell vaccine strategies which target sector residues lying on the p24 intra hexamer interfaces, while avoiding targeting residues on the p24-SP1 interface. In the former case, such targeting seemingly compromises viral fitness by disrupting the formation of stable HIV capsid [24], which appears quite difficult to restore through compensatory mutations. In contrast, restoring fitness costs associated with destabilization of the p24-SP1 interface appears less difficult.

**Fig 6.**
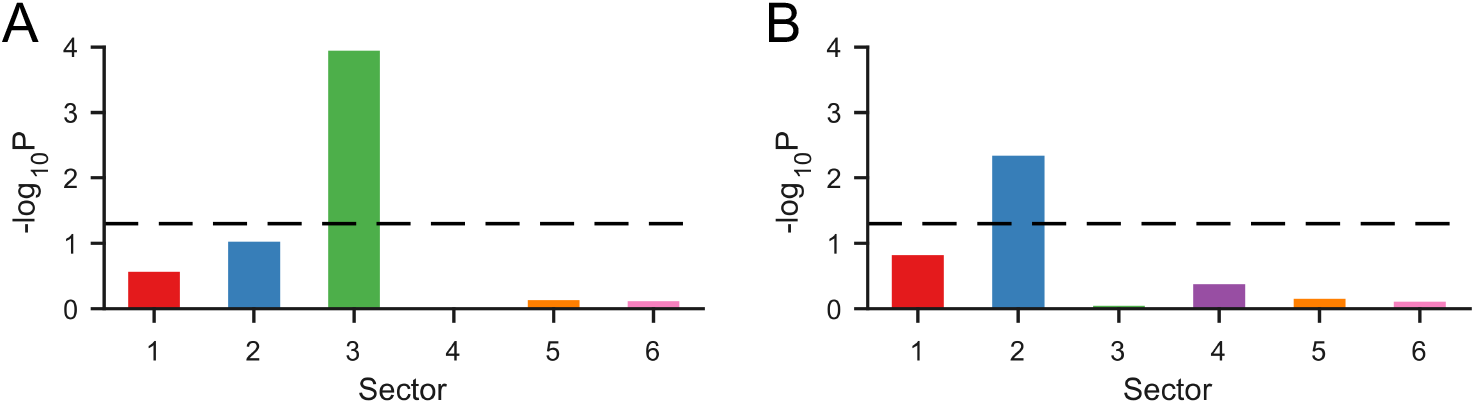
Immunological significance of HIV Gag sectors obtained using RoCA. Association of sectors to (A) viral control and (B) rapid disease progression. The vertical axis of each plot shows the statistical significance of the association, measured as – log_10_ *P*, and the black dashed line corresponds to the conventional threshold of statistical significance, *P* = 0.05; any value above this line is considered statistically significant.

These results were contrasted against a previous analysis of HIV Gag [11], in which an inferred sector based on a classical PCA approach (a slight variant of the approach [12], discussed earlier) was also found to associate with LTNP. Scanning this sector against the Gag biochemical domains (Table 1) revealed a significant association with the p24 intra-hexamer interfaces (as pointed out in [11]), but also with the p24-SP1 interface (S6 Fig). Hence, while reaffirming the importance of targeting interfaces within p24 hexamers, different conclusions were established regarding p24-SP1, suggesting that this interface should be targeted, rather than avoided. This important distinction arises as a consequence of the methodological differences between RoCA and the previous methods [11,12], as discussed previously.

By integrating our observations with population-specific HLA allele and haplotype information, candidate HIV immunogens eliciting potentially robust T cell responses can be proposed [11,12]. A more detailed investigation along these lines, as well as broadening the analysis to other viral proteins, is planned to be carried out in future work.

## Discussion

Characterizing the co-evolutionary interactions employed by HIV and HCV is an important problem. These interactions reflect the mutational pathways used by each virus to maintain fitness while evading host immunity. However, they are not well understood and pose a significant challenge for vaccine development. By applying statistical analysis to the available cross-sectional sequence data, we showed that for multiple HIV/HCV proteins the interaction networks possess notable simplicity, involving mainly distinct and sparse groups of interacting residues, which bear a strikingly modular association with biochemical function and structure. Essential to unraveling this phenomena was the introduction of a robust inference method.

Our approach is particularly well motivated for the “internal” proteins of chronic viruses such as HIV and HCV that are subjected to broadly directed T cell responses. For such proteins, and for HIV in particular, recent experimental and computational work has provided evidence that the population-averaged mutational correlations are reflective of intrinsic interactions governing viral fitness. This was shown to be a consequence of multiple factors which influence the complex evolutionary dynamics of HIV, including the extraordinary diversity of HLA genes in the human population which place selective pressures on diverse regions of the protein, thereby promoting wide exploration of sequence space, in addition to the tendency of mutations to revert upon transmission between hosts [4–6]. An additional important evolutionary factor is that of recombination, which introduces diversity through template switching during viral replication. A consequence of recombination is that it breaks mutational correlations between residues that are distant in the primary structure. That is, higher rates of recombination should lead to shorter-range correlations and vice-versa. This fact is reflected by our inferred co-evolutionary sectors for HIV and HCV. Specifically, the HIV protein sectors are quite localized, with a median separation in the primary structure of up to 140 residues (sector 6 of HIV Gag), while those of the HCV proteins are well separated with a median separation of up to 4S0 residues (sector 1 of HCV NS3-4A). These observations are consistent with the fact that HIV has recombination rates which are substantially higher than those of HCV [46].

In general, the predicted sectors primarily comprise residues within the corresponding biochemical domains and a few other residues which are close in either primary or tertiary structure. However, these sectors also include a small proportion of residues which are distant from those within the respective biochemical domains (S8 Fig) and thus, appear to influence the associated structure or function by an allosteric mechanism. Such long-range interactions have been reported to play a role in maintaining viral fitness and facilitating immune evasion [47–49]. Allosteric interactions have also been observed in the co-evolutionary sectors of different protein families obtained previously with the SCA method [14–17].

The identified sectors for each viral protein together comprise between 35%–60% of the total residues in the protein (Fig 2A and S1 Fig). This is consistent with the sparse sectors of co-evolving residues observed in different protein families using the SCA method [14–17]. One may ask however about the role of non-sector residues, i.e., those not allocated to any sector. Similar to the observations in other proteins [14–17], our analysis suggests that non-sector residues evolve nearly independently, with associated biochemical domains being impacted only by individual mutations at these residues.

While our analysis has focused primarily on viral proteins, the proposed RoCA approach is general and may be applied more broadly, provided that the studied proteins are reasonably conserved. As an example, we computed sectors for the S1A family of serine proteases and compared these with results obtained previously with the SCA algorithm [14,15]. Similar to SCA, RoCA yielded three co-evolutionary sectors which had statistically-significant associations with distinct phenotypic properties; namely thermal stability, enzymic activity, and catalytic specificity (S9 Fig). We point out however that the very notion of a “sector” as defined previously for protein families [14–17] has some conceptual differences to that considered for the viral proteins in this work. Specifically, while for the HIV/HCV proteins, sectors are seen to reflect the mutational pathways employed to facilitate immune escape; for the protein families, they were interpreted quite broadly as representing general features of protein structures that reflect evolutionary histories of conserved biological properties. That is, the concept of a “sector” was defined as extending the classical notion of conservation to incorporate higher order constraints by embracing correlations between protein positions. With this objective in mind, employing a conservation-weighted correlation measure, as specified by SCA, seems appropriate. Nonetheless, the RoCA sectors produced for the serine proteases, based on an unweighted Pearson correlation measure, further attest to the importance of residue interactions in mediating fundamental protein functions.

For the HIV/HCV viral proteins under study, the relation between the biologically important residues (reflected by the biochemical domains) and conservation was not clearly apparent (S10 Fig). In fact, a significant and particularly surprising aspect of our analysis is the substantial extent to which the *correlation patterns, with no regard for conservation*, encode information regarding qualitatively distinct phenotypes including structural units—virus-host and virus-virus protein interactions—and functional domains. The identified sectors may therefore also be seen as *predictors* of important biochemical domains. For each of the four viral proteins under study, there is at least one sector with unknown biochemical significance. Subsequent experimentation, such as mutagenesis experiments targeted at the identified sector residues, could therefore provide new insight which furthers the current understanding of HIV and HCV. Particularly interesting is the poorly understood NS4B protein of HCV, for which any biochemical activity underpinning the leading two sectors—representing the strongest co-evolutionary modes—have yet to be resolved.

## Materials and methods

### Sequence data: Acquisition and pre-processing

The sequence data for HIV-1 clade B Gag and Nef was obtained from the Los Alamos National Laboratory HIV database, http://www.hiv.lanl.gov/. We restricted our analysis to drug-naive sequences and any sequence marked as problematic on the database was excluded. To avoid any patient-bias, only one sequence per patient was selected. After aligning the sequences based on the HXB2 reference, they were converted to a *N* × *M* amino acid multiple sequence alignment (MSA) matrix, where *N* denotes the number of sequences and *M* denotes the number of amino acid sites (residues) in the protein. The downloaded sequences may include a few outliers due to mis-classification (e.g., sequences assigned to an incorrect subtype or clade) in the database. Such outlying sequences were identified and removed using a standard PCA clustering approach. This yielded *N* = 1897 and *N* = 2805 sequences for HIV Gag and Nef, respectively. Moreover, the fully conserved and problematic residues (with blanks or gaps greater than 12.5%) were eliminated, resulting in *M* = 451 variable residues for Gag and *M* = 202 for Nef. Similarly, the sequence data for HCV subtype 1a NS3-4A and NS4B was downloaded from the Los Alamos National Laboratory HCV database, http://www.hcv.lanl.gov/. The downloaded HCV sequences were then aligned based on the H77 reference and converted to the amino acid MSA. Applying the above-mentioned pre-processing resulted in *N* = 2832 sequences for NS3-4A and *N* = 675 sequences for NS4B, with an effective length of *M* = 482 for NS3-4A and *M* = 190 for NS4B.

The processed amino acid MSA matrix **A** = (*A_įj_*) was converted into a binary matrix **B**, with (*i,j*)th entry

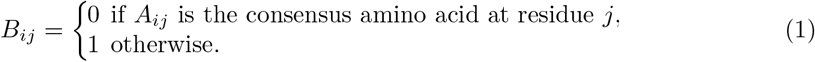

Thus, ‘0’ represents the most prevalent amino acid at a given residue and ‘1’ represents a mutant (substitution). This is a reasonable approximation of the amino acid MSA, given the high conservation of the internal viral proteins under study (S7 Fig).

The binary sequences in **B** are generally corrupted by the so-called *phylogenetic effect*, which represents ancestral correlations. A comparatively large eigenvalue is observed in the associated correlation matrix due to these phylogenetic correlations [11,12]. Following previous ideas [11,12], such effects are reduced using standard linear regression. The resulting data matrix, denoted by 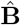, was the base for computing the correlations used to infer sectors. Specifically, we computed the *M* × *M* sample correlation matrix, along with its spectral decomposition, given by

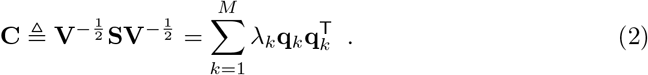

Here, **S** is the sample covariance matrix with entries 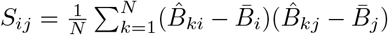 where 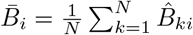 is the sample mean, while **V** is a diagonal matrix containing the sample variances, i.e., *V_ii_* = *S_ii_*, and *λ_k_* and **q**_*k*_ are the *k*th-largest eigenvalue of **C** and its corresponding eigenvector, respectively. The superscript ^**Τ**^ denotes vector transposition.

**Algorithm 1.**
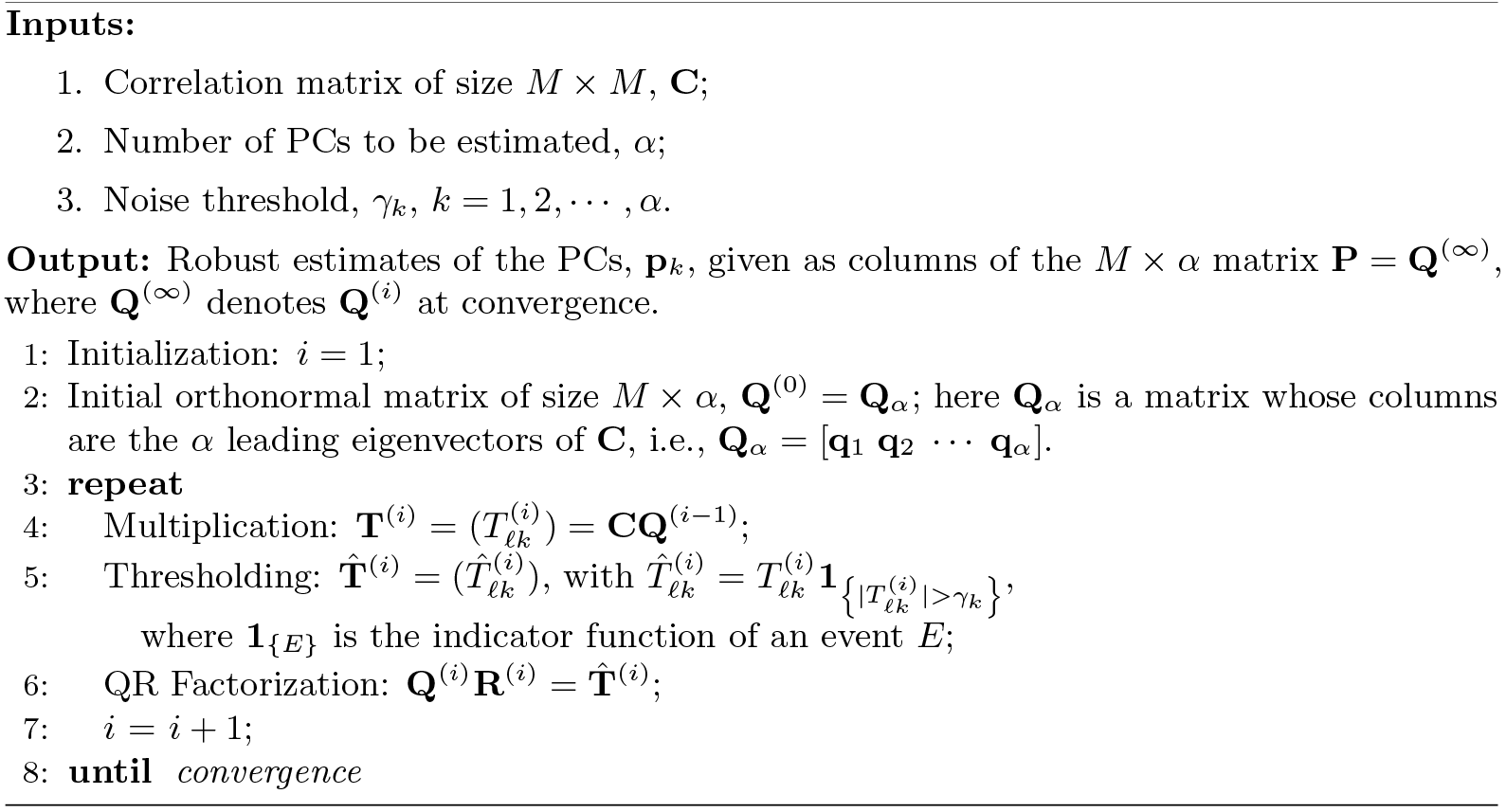
Corr-ITSPCA Method

### RoCA method

We introduced an approach based on robust PCA methods to accurately estimate the PCs (i.e., the leading eigenvectors) of the correlation matrix, which were then directly used to identify sectors. In particular, we considered the iterative thresholding sparse PCA (ITSPCA) method which, in short, is a combination of the standard orthogonal iteration method [50], used to compute the eigenvectors of a given matrix, and an intermediate thresholding step which filters out noise in the estimated PCs. However, the original ITSPCA method was not directly applicable to our correlation-based sectoring problem, since it was designed primarily for covariance matrices, and it involved a variance-dependent coordinate pre-selection algorithm which is no longer suitable. As such, for RoCA, we developed a version (called Corr-ITSPCA, see Algorithm 1) which is appropriately adapted to operate on correlation matrices, and we designed automated methods for tuning the relevant parameters; specifically, the number of significant PCs *α* and the noise threshold *γ*_*k*_.

Such automated design is crucial to obtain accurate results, as these parameters control respectively the number of sectors that we infer and the number of residues included in each sector. Note that this is a rigorous and principled design approach, as opposed to an *ad hoc* approach considered previously by the authors to uncover vaccine targets against the NS5B protein of HCV [51]. These parameters are designed as follows:

### Number of significant PCs, *α*

The design relies on the observed deviations from a null model. Specifically, we generate randomized alignments under the null hypothesis that no genuine correlations are present in the data. This is simply obtained by randomly shuffling the entries of each column of 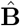, effectively breaking any existing genuine correlation in the data set, and yielding a randomized (null-model) alignment. This shuffling procedure is repeated 10^5^ times and, for each randomized alignment, the maximum eigenvalue of the sample correlation matrix is recorded. The maximum of the recorded eigenvalues, denoted by 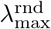, is used as an upper bound of statistical noise so that any eigenvalue of **C** exceeding it is identified as a relevant spectral mode, i.e., as a genuine contribution to the correlation. Thus, the number of significant eigenvectors is set to

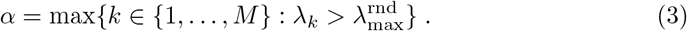

### Noise threshold, *γ_k_*

During the initial iteration, the matrix **T** subject to the thresholding step in Algorithm 1 is given by

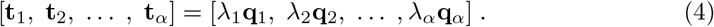

Our design looks for a suitable threshold which eliminates variables that appear uncorrelated (i.e., non-sector variables), for which the corresponding entries of **q_*i*_** correspond purely to sampling noise for every *i* = 1,…,*α*. (Note that Eq (4) only holds for the first iteration of Algorithm 1. However, from [18], coordinates that are set to zero in the first iteration remain zero in subsequent iterations.) Assume that there are *M*_ns_ (unknown) non-sector residues, which we denote by 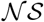, and let 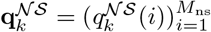 represent **q**_*k*_ but restricted to the coordinates in 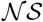 only. The proposed threshold design is based on a statistical description of the worst-case non-sector coordinate; such description relies on the observation that the sample correlation matrices generated with HCV and HIV sequence data have spectral characteristics that are reminiscent of so-called “spiked” correlation models of random matrix theory (Fig 1). To be more specific, we exploit theoretical properties derived for spiked models (Theorems 4 and 6 in [52]) concerning the asymptotic distributions of sample eigenvalues and eigenvectors. Upon particularizing those results to the specific eigenvector structure conveyed by the sectors, they indicate that the coordinates of 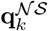 are distributed, up to scaling, as those of a vector that is uniformly distributed on the (*M* – *α*)-dimensional unit sphere. Such a vector is well-known to admit an equivalent representation as a rescaled vector of independent standard Gaussians (i.e., scaled to unit norm). Hence, with *N* and *M* both sufficiently large, *M* ≫ *α*, and *η* = *M/N*, these results along with some basic arguments lead to

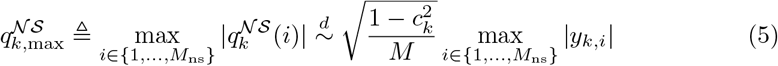

for each *k* = 1, …, *α*, where the *y_k,i_* are independent standard Gaussian random variables, and where the notation 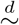 represents “equivalence in distribution”. Here,

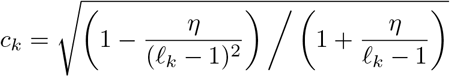

which is a function of the quantity

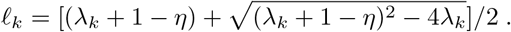

The factor 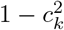 in Eq (5) stems from a complex statistical analysis given in [52], and represents the total statistical noise variance accumulated across all non-sector coordinates 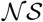 of **q**_*k*_. This quantity increases with increasing *η* = *M/N*, and its value may be quite large, particularly for scenarios in which the number of samples *N* are not substantially greater than the number of protein residues *M* (i.e., the relevant case for our viral data sets). [As a technical aside, we note that there is a condition on the minimum value of 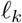 (equivalently *λ_k_*) in order for the above relations to hold (see [52]); though, this condition will typically be obeyed as a consequence of the shuffling procedure used to infer *α*, which selects “sufficiently strong” spectral modes.]

Based on Eq (5), by standard arguments from order statistics, the cumulative distribution function of 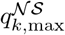 is given by

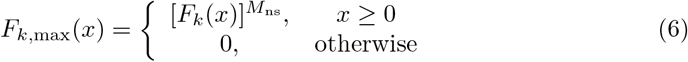

where

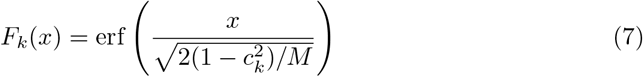

with erf the Gaussian “error function”. Note that all parameters of this distribution are functions of observable quantities (e.g., *λ_k_, M*, and *N*), with the exception of the number of non-sector coordinates, *M*_ns_. To account for this, we invoke a worst-case assumption, replacing *M*_ns_ with its upper bound, *M*. We may then set a threshold for the coordinates of **q***_k_* based on this distribution, considering a suitable percentile. Here, taking a 95% cut-off,

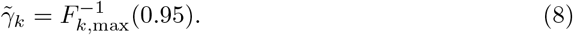

Numerical simulations (S2 Text) demonstrate that this choice of 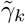 serves as a good compromise between the ability to accurately capture the sector residues of interest (i.e., a high true positive rate), while rejecting most of non-sector residues (i.e., a low false discovery rate). Finally, since the threshold is applied to the column vectors **t**_*k*_ (Eq (4)), rather than the **q**_*k*_, the noise threshold *γ_k_* is chosen as

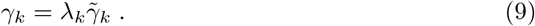

The iterative procedure in the Corr-ITSPCA method (Algorithm 1) is similar to the standard orthogonal iteration procedure used to obtain the eigenvectors of a matrix [50]. However, addition of the intermediate thresholding step helps to identify a subspace spanned by the significant PCs such that there is no contribution from the non-sector residues in the estimated PCs. Moreover, in the process of obtaining an orthogonal subspace, this iterative procedure accurately infers the coordinates contributing to each PC by resolving spurious overlap between the support (non-zero components) of all the significant PCs, as demonstrated by ground-truth simulations (S11 Fig).

From the sample correlation matrix **C**, the robust estimate of the PCs was obtained using the Corr-ITSPCA method (Algorithm 1), with *α* and *γ_k_* designed as above. The estimated PCs **p**_*k*_, *k* = 1, …, *α*, produced by Algorithm 1, were then used to form *α* sectors as

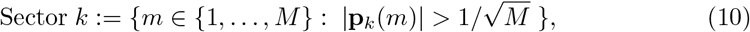

where |**p**_*k*_ (*m*)| is the absolute value of the *m*th coordinate of **p**_*k*_. Note that the estimated **p**_*k*_ do not generally have strict zero entries in the non-sector coordinates, but may contain very small values due to residual noise. As such, a small threshold was applied (Eq (10)) to form sectors in an automated way. The spurious entries were generally quite distinguishable from the relevant coordinates, even from simple visual inspection of **p**_*k*_ (Fig 2A and S1 Fig).

As mentioned above, all fully conserved residues in the MSA were initially excluded from our analysis, as the Pearson correlation involving these residues was not defined. Given the lack of information regarding their potential interactions with other residues, and considering the tendency of neighboring residues in the primary structure to interact with each other, any fully conserved residue in the immediate neighborhood of a sector residue was incorporated into that sector.

### Heat map of cleaned correlations for visualization

In figures 2 and 5, we used heat maps to illustrate the computed sample correlation matrix **C**. As discussed above, the sample correlations were generally corrupted by statistical noise due to the finite number of available sequences. Thus, for a better visualization and, in particular, to appreciate the strong correlations within the inferred sectors, the sample correlation matrix was cleaned from statistical noise by thresholding the sample eigenvalues in such a way that the significant *α* spectral modes (Eq (3)) were kept unaltered, while the remaining eigenvalues (which do not appear to contribute genuine correlations) were collapsed to a constant. Specifically, the cleaned sample correlation matrix was obtained as

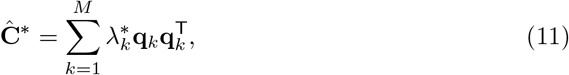

where

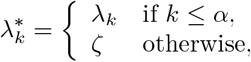

with ζ a constant value such that the trace of 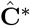 remained normalized (equal to *M*). Note that 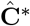 is not a standard correlation matrix as 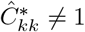. A standardized version was then computed as

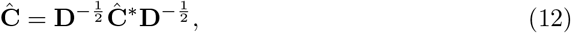

where **D** is a diagonal matrix with 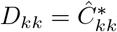, and used to depict the cleaned correlations as a heat map.

### Statistical independence of inferred sectors

We introduced a metric called “normalized entropy deviation (NED)” to quantify the extent to which two groups of residues are statistically independent of each other. The NED between two sectors *i* and *j* is defined as

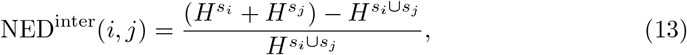

where *s*_*i*_ is a set comprising the five residues with largest correlation magnitude of sector *i* and *H^s_i_^* the entropy of *s_i_* computed from the binary MSA matrix. Specifically, this entropy is computed over all *κ* = 1, 2, · · ·, 2^#(*s_i_*)^ combinations of the residues in set *s_i_* as follows

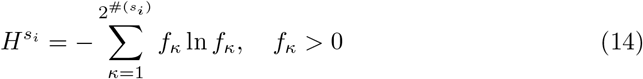

where *f_κ_* is the frequency of the combination *κ* in the MSA and #(*s_i_*) is the cardinality of set *s*_*i*_. In theory, if two given sectors are perfectly independent, the sum of the entropies of the individual sectors must be equal to the entropy of both sectors taken together, resulting in NED^inter^ = 0. In practice however, a small non-zero value of NED^inter^ is expected due to finite-sampling noise, even if the sectors are independent. We obtain an estimate of it by constructing a null case, where the entries of the MSA corresponding to the sets *s_i_* and *s_j_* are randomly shuffled in such a way that any correlation between the two sets is essentially eliminated, while the correlations between residues in an individual set remain unaltered. Using Eq (13), NED^inter^ is computed for 500 such randomly shuffled realizations of the MSA and the average value (referred to as NED^random^) represents the null (lower) reference value for NED^inter^ which is expected if the two sectors are independent. Substantial deviations from NED^random^ should reflect correlation between the sectors. In order to quantify the extent of such deviations in a clearly correlated case, we computed an upper reference NED^intra^, obtained when the residues in both sets *s_i_* and *s_j_* belong to the same sector. It is defined as

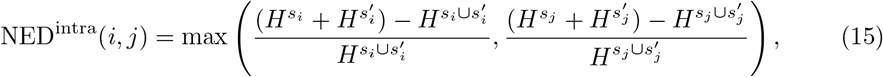

where 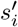 is the set comprising the five residues with largest correlation magnitude of sector *i* with the residues in *s_i_* excluded.

### Clinical data used in the immunological study

The HLA alleles associated with control and progression of HIV Gag were reported in [53]. A list of Gag epitopes associated with either control or progression was compiled using the data from the Los Alamos HIV Molecular Immunology Database (http://www.hiv.lanl.gov/content/immunology) and is presented in S1 Table.

### Data and code availability

Accession numbers of all the sequences used in this work are provided in S3 File. Source code for the proposed RoCA method along with the code for reproducing all figures is available at https://github.com/ahmedaq/RoCA.

## Supporting information

**S1 Text. Details of the PCA inference method [11,12].**

**S2 Text. Simulation study: Statistical robustness of RoCA.**

**S3 Text. Analysis of the SCA method.**

**S1 Fig. Co-evolutionary sectors revealed by RoCA for (A) HIV Nef, (B) HCV NS3-4A, and (C) HCV NS4B proteins**. The first row of each panel shows the biplots of the estimated sparse PCs that are used to form RoCA sectors. The sector residues are represented by circles according to the specified color scheme, while overlapping residues (belonging to more than one sector) and non-sector (independent) residues are represented as gray and white circles, respectively. The heat map of the cleaned correlation matrix, with rows and columns ordered according to the residues in the RoCA sectors, shows that the sectors are notably sparse and uncorrelated to each other. The second row of each panel plots the location of RoCA sector residues in the primary structure. The sector residues are colored according to the specifications in Fig 2A. The third row of each panel shows the statistical independence of the RoCA sectors, quantified using the normalized entropy deviation metric.

**S2 Fig. Robustness of the association of HIV Gag sectors 3 and 6 to the respective structural (interface) domains for different values of the contact-defining distance** *d* **between the alpha-carbon atoms.** The protein chains in all structures are shown in gray. The structurally important residues (i.e., interface residues) present in a sector are shown as red spheres and the remaining residues of the interface are shown as blue spheres. The first, second, and third column show the interface residues obtained with *d* = 7, 8, and 9Å, respectively. Crystal structure of (A) the p24 hexamer (PDB ID 3GV2), (B) the p24 pentamer (PDB ID 3P05), and (C) the p24 inter-hexamer interface (PDB ID 2KOD). All the associated results in the main text were shown for *d* = 7Å. The association of sector 3 and sector 6 to the corresponding interfaces remains statistically significant for different values of *d*.

**S3 Fig. Comparison of the PCA-based method [12] and the proposed RoCA method.** (A) Comparison of sector sizes obtained using PCA and RoCA for the studied viral proteins. The y-axis shows the number of residues present in each sector. (B) Associations of sectors produced by the PCA-based method [12] with the biochemical domains of the studied viral proteins. Only the sectors having statistically significant association with any biochemical domain are presented. The sectors are colored according to the scheme in Fig 2A. The area of each bubble reflects the statistical significance of the associated result, measured as − 1/ log_10_ *P*, and the black circles indicate the conventional threshold of statistical significance, *P* = 0.05; any *P*-value lower than that is considered statistically significant. The *P*-values associated with non-significant associations (*P* > 0.05) are displayed inside the black circle. (C) Comparison of the robustness of RoCA and PCA [12] methods to finite sampling using binary synthetic data. The parameters used in the simulation were *M* = 500 residues and *r* = 5 non-overlapping units *S_i_* of size 12%, 10%, 8%, 6%, and 4% of *M*, respectively for *i* = 1, …, 5. The corresponding 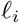 were set to equally spaced values between 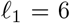 and 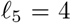, and to model the phylogenetic effect, we set 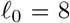. To test the finite-sampling effect, results are presented for varying number of samples *N* = 1000, 2000, and 4000 corresponding to 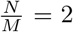, 4, and 8, respectively. The sectors inferred using RoCA and PCA [12] are compared using mean TPR, mean FDR, and the maximum percentage mismatch PM_max_. In each box plot, the black circle indicates the median, the edges of the box represent the first and third quartiles, and whiskers extend to span a 1.5 inter-quartile range from the edges.

**S4 Fig. Sectors revealed by the PCA-based method [12] for (A) HIV Nef, (B) HCV NS3-4A, and (C) HCV NS4B proteins.** The first column displays the bar plots showing the merging of multiple RoCA sectors in the sectors revealed by the method in [12]. The vertical axis of each plot shows the percentage of residues within the different RoCA sectors that fall into the prescribed sector. The second column shows the biplots of the PCs which are post-processed to form sectors in [12]. The sector residues are represented by circles according to the specified color scheme, while independent (non-sector) residues are represented as white circles.

**S5 Fig. Comparison with other co-evolutionary methods**. (A) Comparison of the biochemical association of the sectors inferred by the three sectoring methods: 1) The proposed RoCA method, 2) the PCA method [12], and the SCA method [14]. (B) Biochemical association of HIV Gag sectors inferred using alternative co-evolution methods available in the literature (reviewed in [13]). Specifically, the inferred sectors were based on: 1) Direct coupling analysis (DCA) [54], 2) a mutual information (MI) based method [55], 3) McLachlan based substitution correlation (McBASC) method [56], 4) observed minus expected square (OMES) method [57], and 5) evolutionary trace (ET) method [58]. The MI and DCA methods were implemented using the code provided in [54]. For a fair comparison, the similarity-based sequence weighting of DCA was not applied; however, a pseudo-count value of 0.5 was used (as specified in [54]) to avoid singularity issues during inversion of the covariance matrix in DCA. The McBASC [56] and OMES [57] methods were implemented using the code provided in [59]. The ET method was run from the web-based server provided at http://mammoth.bcm.tmc.edu/ETserver.html. None of these methods, except the ET method, were originally designed to explicitly produce *sectors* of co-evolving residues, but to simply assign a score to each pair of residues in the protein, with a high score indicating a high probability of the associated pair to be in contact in the tertiary structure. Nonetheless, we formed a single sector based on these pairwise scores. This was done by aggregating those pairs of residues deemed to be *significantly* interacting, corresponding to pairs with an associated score larger than *β* = 2 standard deviations above the mean of the overall distribution of scores. Different choices of *β* yielded qualitatively similar results (not shown). The ET method combines information of the cross-sectional conservation (single-residue conservation in the MSA) and the conserved residues in different branches of the phylogenetic tree (associated with the input MSA) to assign a score to each protein residue. In this algorithm, a lower score reflects higher importance of the residue. Thus, we formed a sector by including those 20% of residues with the lowest scores (as mentioned in [58]). The sector predicted by these methods, except the ET method, showed no statistically significant association to any biochemical domain in HIV Gag. The sector predicted by the ET method was found to be associated with the P7-Zinc-Finger domain. Note that we also tested the multiple correspondence analysis (MCA) based S3det co-evolution method [60] using the web-based server provided at http://treedetv2.bioinfo.cnio.es/treedet/index.html. However, no results could be obtained due to its high computational complexity when applied to the (large) Gag protein. (C) Biplots of all possible pairs of the top six PCs—after discarding the leading eigenvector representing the phylogenetic effect—of the SCA matrix, used to form HIV Gag sectors with SCA. Sector residues, overlapping residues, and non-sector residues are represented with the same color scheme of Fig 2A.

**S6 Fig. Biochemical association of the HIV Gag sectors reported in [11]**. For all structural interfaces, the contact-defining distance between the alpha-carbon atoms was set to *d* = 7Å. The inference method in [11] is similar to the PCA approach [12] tested in this paper, mainly differing in the procedure to form sectors from eigenvectors; specifically, the method in [11] formed sectors from visual inspection of eigenvector biplots, whereas an automated procedure was applied in [12]. Similar to [12], the method in [11] produced relatively large sectors that did not show the highly resolved and modular biochemical association revealed by RoCA (Fig 3). Importantly, no sector reported in [11] was found to be associated with rapid progression to AIDS. This was due to the fact that the residues in the biochemical domain important in this case (p24-SP1 interface; see *Results*) were mixed up with the control-associated residues in sector 3.

**S7 Fig. Comparison of the entropy per residue of the amino acid MSA and that of the binarized MSA for each studied viral protein.** A high positive Pearson correlation *r* (close to 1) with very high statistical significance (very small *P*) is obtained between the entropy per residue of the amino acid MSA (H_*i*_) and that of the binarized MSA 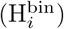 in all cases; for details on the entropy computation, see [12]. This demonstrates that the binary MSA is a good approximation of the amino acid MSA for the considered viral proteins.

**S8 Fig. Presence of long-range interactions in the predicted sectors.** Circular plots showing both the short and long-range mutational interactions in the six sectors predicted for HIV Gag. The interactions among residues in a sector are represented with colored lines following the scheme specified in Fig 2A. For better visualization, only the strong interactions (|*Ĉ_ij_*| > 0.1) involved in each sector are shown.

**S9 Fig. Biochemical association of the RoCA sectors obtained for the S1A serine protease family, in comparison with the SCA sectors produced by [14].** The RoCA sectors were inferred from the same sequence data used by [14]. The relevant biochemical domains in this case were identified as distinct sets of residues involved with thermal stability [61,62], basic chemical function of this enzyme family, and catalytic specificity [63, 64]. An important distinction here with respect to the HIV and HCV proteins is that the leading eigenvector obtained in the estimation of the SCA and RoCA sectors was associated with two biochemical functions; thermal stability and enzymic activity. In [14], two sub-sectors were formed from this eigenvector based on the sign of its elements; specifically, the red sub-sector was formed using the positive elements, while the blue sub-sector was formed using the negative elements. The same strategy was applied for RoCA, resulting in the corresponding sector 1 being divided into two sub-sectors, 1a and 1b.

**S10 Fig. Analyis of the conservation of the residues within the biochemical domains of each studied viral protein.** We categorize the residues in each protein into fully conserved and variable residues. The variable residues are further divided into five conservation quantiles Q1, Q2, …, Q5, with quantile Q1 consisting of the 20% most-conserved residues, Q2 the next 20% most-conserved residues, and so on. The y-axis represents the fraction of biochemically important residues present in each conservation group for each protein. This result shows that the residues within the biochemical domains of each studied viral protein are well-mixed with respect to conservation.

**S11 Fig. Effect of Corr-ITSPCA iterations on the RoCA sector inference using binary synthetic data**. The parameters used in the simulation study (see S2 Text for details) were *M* = 500 residues and *r* = 5 non-overlapping units *S_i_* of size 12%, 10%, 8%, 6%, and 4% of *M*, respectively for *i* = 1, …, 5. The corresponding 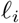were set to equally spaced values between 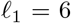 and 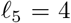, and to model the phylogenetic effect, we set 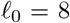. The ratio of the number of samples to the number of residues was fixed at 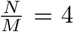 and the simulation was run for 500 Monte Carlo realizations. (A) The maximum percentage mismatch PM_max_ between sectors formed using the PCs estimated at a particular Corr-ITSPCA iteration and the corresponding units. PM_max_ decreases as the number of iterations increases, demonstrating that the iterative procedure in Corr-ITSPCA helps to accurately predict the true units. Here, the intermediate iteration corresponds to half of the total number of iterations Corr-ITSPCA took to converge in each Monte Carlo realization. (B-D) Illustration of the convergence of RoCA sectors to the corresponding units in a single Monte Carlo realization for the simulation setting of (A). Snapshot of (B) PCs representing the true units (here, the first PC represents phylogeny while the subsequent five PCs represent the five units), (C) PCs of the sample correlation matrix (constructed using the phylogeny-filtered MSA) used in forming PCA sectors, and (D) PCs at first, fourth, and eight (last) iteration of the Corr-ITSPCA method.

**S1 Table. List of HLA class I restricted epitopes associated with long-term non-progressors (LTNP) and rapid progressors (RP) in HIV Gag.**

**S1 File Detailed list of experimentally-identified biochemical domains of the studied viral proteins.**

**S2 File. List of residues in the sectors inferred using RoCA for all four viral proteins.**

**S3 File. List of accession numbers of all the protein sequences.**

## Acknowledgments

We are particularly grateful to Arup Chakraborty for extensive discussions over the duration of this work. We also thank Iain Johnstone, Karthik Shekhar, John Barton, and Raymond Louie for providing useful input on an earlier version of the manuscript.

## Author Contributions

A.A.Q., D.M.J. and M.R.M. all contributed to designing the research, developing the computational techniques, analyzing and interpreting the results, and writing the paper. A.A.Q. performed the computations involving the sequence data and compiled from the literature the existing biochemical information of the proteins under study.

